# Signal and noise in circRNA translation

**DOI:** 10.1101/2020.12.10.418848

**Authors:** TB Hansen

## Abstract

Within recent years, circular RNAs (circRNAs) have been an attractive new field of research in RNA biology and disease. Consequently, numerous studies have been published towards the disclosure of circRNA biogenesis and function. Initially, circRNAs were described as a subclass of cytoplasmic non-coding RNA, however, a few recent observations have proposed that circRNAs may instead be templates for protein production. The extent to which this is the case is currently debated, and therefore using rigorous data analysis and proper experimental setups is instrumental to settle the current controversies. Here, the conventional experiments used for detecting circRNA translation are outlined, and guidelines to distinguish signal from the inherent noise are discussed. While these guidelines are specific for circRNA translation, most also apply to all other aspects of non-canonical translation.

## Introduction

Within recent years, circRNAs have emerged as a fascinating new class of molecules that based on their closed covalent structure exhibit high stability in cells [1,2]. CircRNAs have been identified in almost all studied eukaryotes, and the expression has been shown to be highly tissue specific [3,4]. While the landscape and expression profiles of circRNAs have been examined in detail, the functional relevance of circRNAs is still largely unsettled. Initially, spawned by the discovery and characterization of ciRS-7/cdr1as [5–7], many circRNAs are proposed as putative miRNA regulators despite stoichiometric challenges [8,9] and absence of miRNA response element enrichment in circRNA sequences [10,11]. Recently, a subset of circRNAs were published to show the ability to encode circRNA-specific peptides [12–14]. In most cases, the circRNAs share the translation initiation site with the corresponding host-gene, and thus the resulting peptides are predicted to resemble the very N-terminal region of the mRNA-derived protein. To identify and study circRNA-derived translation, different approaches have been used, including ectopic circRNA expression, polysome gradients, ribosome footprinting (RiboSeq), and mass-spec analysis. Here, the common pitfalls using these approaches are discussed along with requirements for claiming translation.

## Results

### Ectopic expression

The most convenient approach to study circRNAs translation is to device an overexpression vector, transfect cell-line of interest, and detect protein production using western blot analysis. Even without commercially available antibodies, the overexpression vector can include a tag immediately upstream of the circRNAs-specific stop-codon to detect specifically the circRNAs-derived proteins. However, importantly, all experiments contain unforeseen biases, aberrant outputs, off-targets, technical and biological variation, collectively referred to here as noise. Similarly, for overexpression-based setups, the intended vector-derived output (and its cellular consequences) is presumably only a fraction of noise-derived effects and that is why proper control experiments are essential. Control experiments are required to disentangle signal from noise and to determine the isolated effects of the intended perturbation. For circRNAs overexpression, the control experiment is typically an empty vector, and consequently everything produced from the circRNA expression vector is categorized as signal and circRNA-derived. Unfortunately, to our knowledge, no vector-based overexpression system is sufficiently clean to dismiss any contribution from linear artefacts. Indeed, we and others have shown [11,15], that upon overexpression, the specific sequence produced by back-splicing also occurs in linear concatemers, originating either from so-called ‘rolling circle’ plasmid transcription or non-canonical trans-splicing (Fig. 1A). Here, all features assumed to be circRNA specific, namely the unique back-splicing region, is reconstructed in a linear context, and similarly, any open reading frames (ORFs) defined as circRNAs-specific are now contained in a capped and poly-adenylated mRNA transcript. The problem is not the vector-design used for overexpression, it is the poorly devised control experiment. As previously suggested [11,15], one improved alternative is instead to use vectors containing the circRNAs exon(s) of interest, but lacking the flanking regions required for circRNAs biogenesis (Fig. 1B) to unequivocally detect the circRNAs-derived proteins. Here, aberrant linear exon-repeats are likely unaffected by impaired back-splicing and thus the observed difference between the circRNA expression vector and the control is now attributable as a circRNAs-specific phenotype.

**Figure 1.**
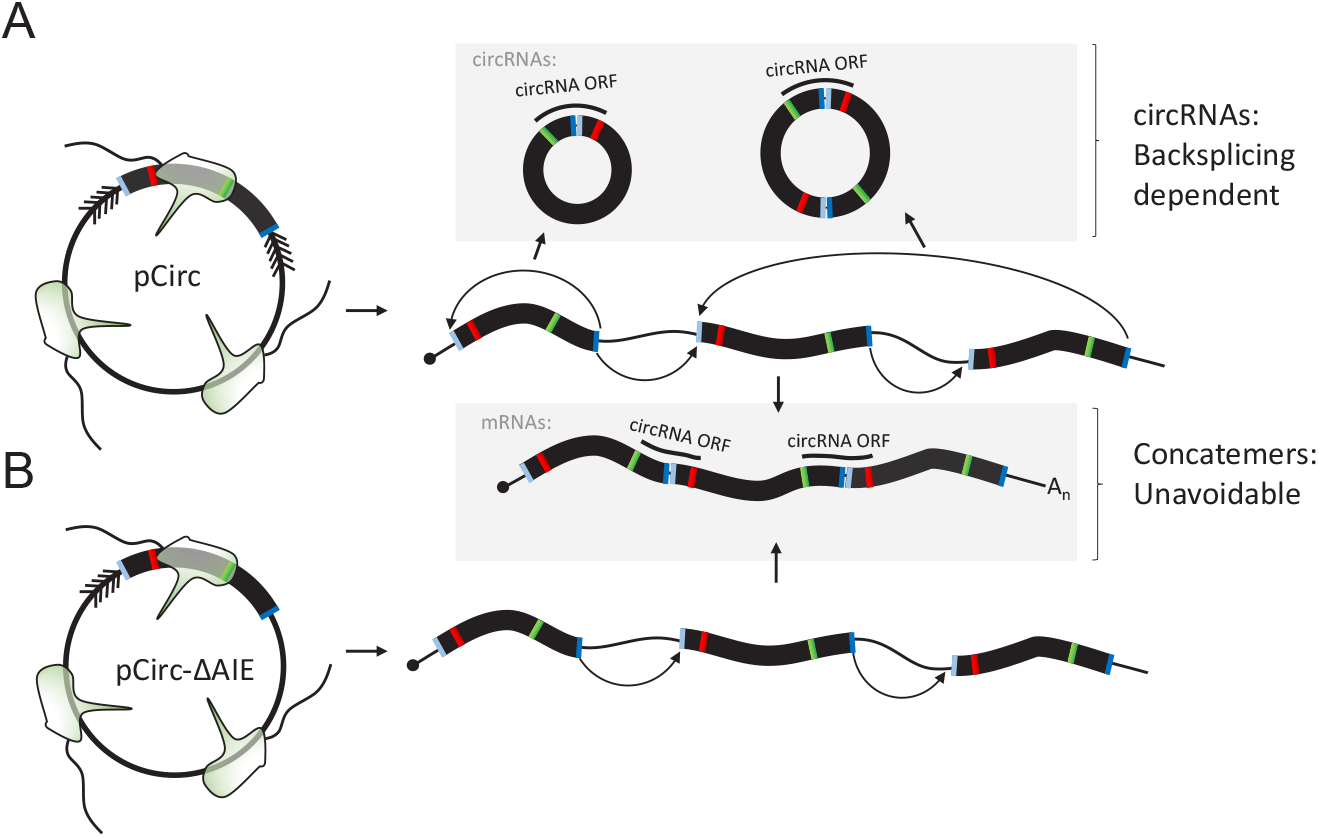
Ectopic circRNAs expression. **A)** Schematic showing the predicted output from a hypothesized ectopic circRNA overexpression vector (pCirc). CircRNA production by backsplicing results in nonlinear exon assembly with the circRNAs specific back-splicing junction and the associated circRNAs-specific peptide sequence (upper scenario, grey box). Rolling circle plasmid transcript results in exon-concatemers producing the non-linear exon assembly in a linear context and the associated protein output is indistinguishable from the circRNAs-derived output (lower scenario, grey box). **B)** Schematic showing the proposed control setup, where the downstream artificial inverted element (AIE) has been removed (pCirc-ΔAIE). Here, backsplicing and circRNA production is impaired and the only relevant output is the linear exon concatemers (lower scenario, grey box).

### Polysome profiling, ribotag and RiboSeq technologies

Classical approaches toward translatome disclosure is to purify ribosomes and quantify the associated mRNA. One embodiment of this approach is the polysome profiling by sucrose gradient fractionation, whereby polysomes are separated from ribosomal subunits and monosomes, and the fractional co-elution of translated mRNA species serves as a proxy for ribosome occupancy on ORFs and thus the level of translation [16]. Another embodiment is the ribotag technique, where a ribosomal subunit, typically RPL22, is tagged allowing full ribosome immunoprecipitation. Here, clever genetic engineering and tissue-specific Cre-recombinase expression, allows for the interrogation of cell-type-specific translation in mouse models [17]. While polysome gradients and ribotag methods greatly enrich for translated RNA species, they do not provide a clear demarcation between the coding and non-coding landscape within cells; some transcripts are subjected to low-level translation while others may associated with the ribosome unspecifically or indirectly (the noise). When using polysome gradients or ribotag qualitatively for circRNA translation, it is imperative to pursue a mutational approach whereby imperative elements, such as the AUG start codon, are disrupted to allow a signal to noise estimate. With current genome editing technologies, it has become possible to engineer specifically such mutations endogenously, although successful identification of homozygous mutants is a laborious approach. And, in most cases, the circRNA and host mRNA ORFs overlap disqualifying a circRNA-specific mutational analysis. Consequently, the urge to utilize vector-based expression for these analyses may prevail, although, again, this requires the ability to separate bona-fide circRNAs from exon concatamers (as discussed in the previous paragraph) using either rigorous RNaseR treatment or northern blotting. In any case, the obtained output should be compared with a translation incompetent mutant to highlight that the ribosome association is indeed coupled to translation.

More recently, high-throughput ribosome foot-printing, coined RiboSeq analysis, has emerged as a powerful technique to discover ribosome-bound open reading frames globally and to quantify translation efficiencies [18,19]. In contrast to polysome gradient and ribotag approaches, RiboSeq only captures ribosome-protected fragments, and thus to demarcate between circRNA and overlapping mRNA transcripts, only back-splice junction spanning fragments are unequivocally derived from circRNA. Like most other experiments, RiboSeq has inherent noise, and inability or failure to estimate signal:noise ratios is potentially a severe problem, particularly when conducting a high-throughput data analysis. A hallmark of active translation is the triplet periodicity or phasing of reads across the ORF. However, capturing the exact phasing of 25-35 nucleotides (nts) RiboSeq reads across the confined circRNAs-specific back-splice junction (BSJ) has obvious limitations, which reduces signal and power significantly. Moreover, attempts to enhance signal by combining the signal from multiple circRNAs has failed to detect RiboSeq phasing [11]. Of note, combining putative ORFs across the back-splice junction of circRNAs pre-assumes the frame of translation, and arguably, this may not necessarily adhere to the same coding frame of the host gene. Then, the alternative approach is to simply count all reads mapping across the back-splice junction irrespective of phasing as a proxy for translation. Here, albeit with general low coverage compared to genuine ORFs, thousands of reads map uniquely to circRNA back-splice junctions (Fig. 2A). Ideally, this suggests that most circRNAs undergo translation but with low rate. The difficult question is then, how many of these reads derive from translating ribosomes and how many are background noise? In an attempt to answer that question, one may assume that the signal:noise ratio (the dataset quality) correlates with the ability to capture phasing across bona fide coding sequences (CDS). Evidently, with this approach, RiboSeq datasets vary dramatically in quality as exemplified by P-site distribution across *β-ACTIN* using high quality and low quality datasets, respectively (Fig. 2B-C). Alarmingly, using the same datasets to quantify the total number of reads on the *β-ACTIN* CDS and across the back-splice junction of circRNAs, there is clearly a tendency towards low quality datasets containing increased number of BSJ-spanning reads, and vice versa for *β-ACTIN* (Fig. 2D). In fact, globally, when contrasting the fraction of reads (reads per mapped million, RPMM) mapping on annotated CDSs with dataset quality (i.e. level of observed phasing), a clear positive correlation is observed. In contrast, the fraction of circRNA mapping reads show negative correlation for all possible read-lengths. In most analyses, the counterfactual scenario, also referred to as the negative control, is an important measure to demarcate signal from noise in high-throughput analysis. Thus, if the definition of signal is the mere presence of reads, the counterfactual scenario should be devoid of reads. Here, with small RNAs, miRNA and snoRNAs, representing a counterfactual scenario, extensive RiboSeq coverage is still observed, but, consistently, similar to circRNAs, the fraction of mi/snoRNAs aligned reads drops subtly with increased RiboSeq quality (Fig. 2E). While circRNAs, miRNAs and snoRNAs in principle could serve as templates for protein production, assuming some level of noise potentially allows detection of all expressed RNA species, and consequently using detection as a proxy for translation fails to appreciate the entire reason for conducting RiboSeq analysis in the first place. Therefore, while reads across circRNAs backsplice junction may at first seem as evidence for circRNAs translation, without detectable phasing the parsimonious explanation is likely a noisy dataset.

**Figure 2.**
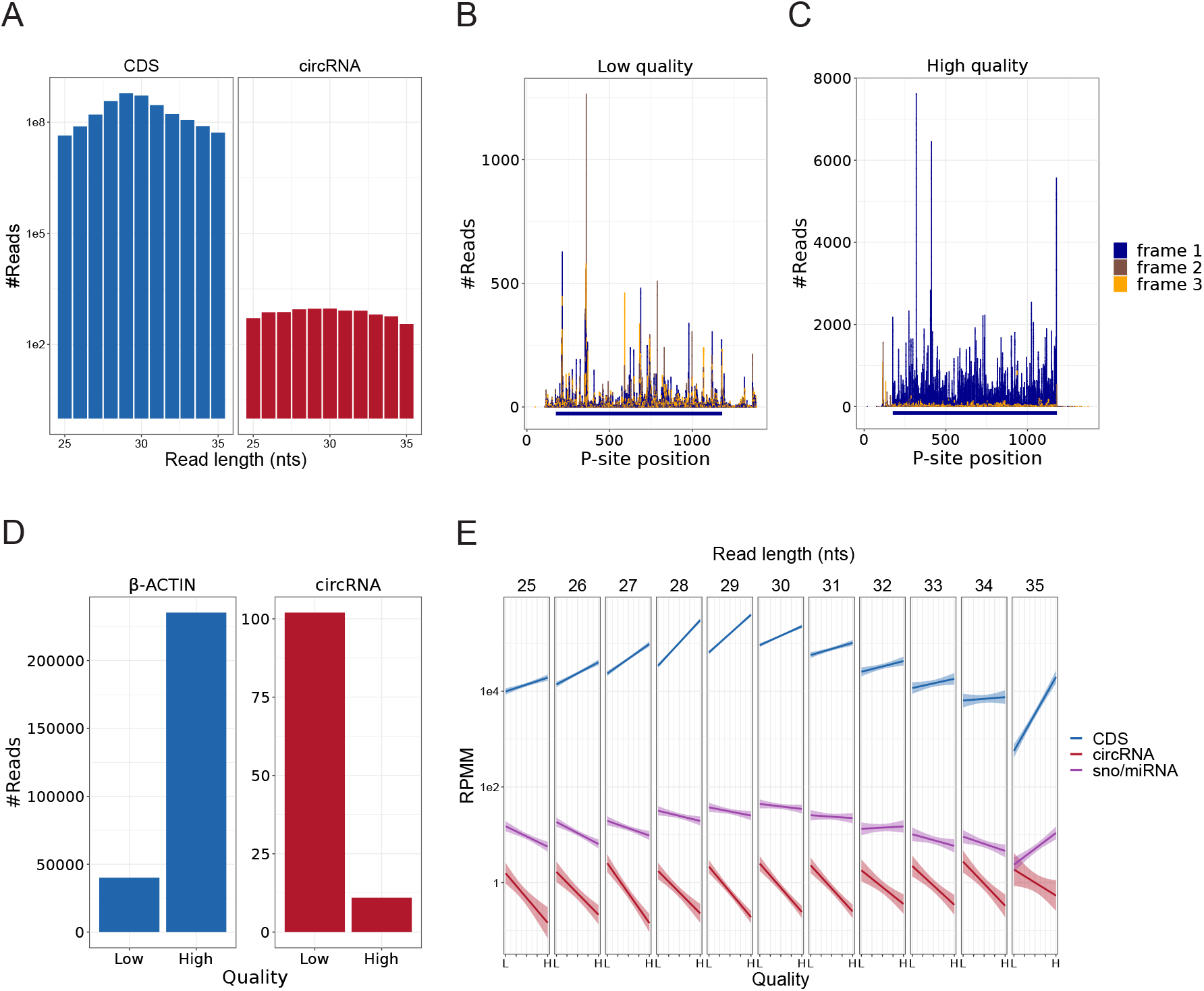
Riboseq analysis. **A)** Read counts from riboseq dataset mapping on the coding sequence (CDS) of *B-ACTIN* (blue, left) or across the backsplice junction (BSJ) of the top1000 expressed circRNAs across the ENCODE RNAseq datasets (red, right) stratified by read length. **B-C)** Examples of 29 nt RiboSeq read distribution across the *β-ACTIN* CDS color-coded by frame (dark blue: on-frame P-sites, brown and orange: off-frame P-sites) for high-quality datasets (top 5%, B) showing clear triplet phasing (enrichment of on-frame reads), and low quality datasets (bottom 5%, C) where no phasing is evident. The horizontal dark blue bar represents the annotated CDS. **D)** As in A, but stratified by dataset quality as in B-C. **E)** Smoothed correlation between ranked dataset quality and fraction of reads (reads per mapped million, RPMM) found on all annotated CDSs (blue), all annotated miRNA and snoRNA genes (not overlapping CDSs, purple), or across the BSJ of the top1000 expressed circRNAs (red), stratified by read length as denoted.

### Mass-spectrometry

In the RNA research field, mass-spectrometry is typically considered a bulletproof approach towards peptide identification. But, unlike most RNA-based techniques, mass-spec analysis has an in-built noise estimator to compute error and false discovery rates (FDR), the so-called target-decoy search strategy [20,21]: All identified peptides have an associated score that signifies the degree of match between the observed and predicted spectrum. Then, to estimate noise, mass-spec analysis is in parallel performed on a positive list of peptide sequences (typically the annotated proteome) and a similar sized list of decoys (often the reverse peptide sequences to preserve amino-acid composition), i.e. the decoys are directly derived from the positive target list. Then, simply put, using the ratio between target and decoy for any given score, the probability of a false assignment is deduced, resulting (when using Maxquant) in a posterior error probability value (PEP). Additionally, by accumulating the number of detected decoys in a ranked list, the overall FDR, i.e. how many false discoveries are expected in the resulting peptides, is available for any particular PEP/score cutoff. While this is very simplified account of the mass-spec statistics, the imperative point is that the final FDR reflects the entire group of peptides identified, and obviously, each individual peptide is not called with equal confidence (PEP). With this in mind, taking a non-random subsample from the total bulk of peptides, the initial FDR does not apply anymore. For instance, extracting the top 5% or the bottom 5% based on PEP values, the signal-to-noise ratio in each subset is dramatically different, and thus the FDR must be re-calculated based on the ratio of positive and decoy peptides in each subsample.

To exemplify the importance of FDR-recalculation, a re-analysis of the circRNA peptides recently published in *Cell* by van Heesch *et al*, 2019 [22] was performed. Again, like the RiboSeq analysis, a counterfactual approach was included that surely should be devoid of signal, here the COVID-19 proteome. Similar to van Heesch *et al*, Maxquant mass-spec analysis was performed on an extensive datasets from human heart [23]. Here, almost 4 million uniquely assigned peptides were derived from the Uniprot proteome assemble, whereas 38 peptides where found as circRNA specific of which 14 span the backsplice junction (Fig. 3A). In addition, Maxquant identifies 79 total COVID-19 peptides, however, as seen for both circRNA and COVID19-derived peptides, the number of matched decoys exceed the number of positive targets (Fig. 3A). Also, importantly, the distribution of PEP values obtained from circRNAs, COVID-19 and the Uniprot proteome are very different (Fig. 3B). Here, the Uniprot signal greatly diverge from noise when PEP is small, whereas the distribution looks very similar for circRNA and COVID19. First of all, this shows that circRNA and COVID19 peptides are a non-random subset of the bulk output, and second, that the signals obtained from circRNAs and COVID19 are inseparable from noise. Consistently, when re-computing the FDR based on the level of circRNA- and COVID-derived decoys, the FDR increases to 0.67 and 0.65, respectively. In fact, not a single possible PEP-value cutoff results in any significant circRNAs-derived peptides using FDR < 0.05 (Fig. 3C), however, alarmingly, four distinct COVID19-specific peptides are called as significant using a PEP < 0.02 cutoff. This, surely, must reflect either insufficient FDR control by the target-decoy strategy (discussed below), or that the detection of only four peptides is very likely by change, which prompts additional caution when only few peptides are identified.

**Figure 3.**
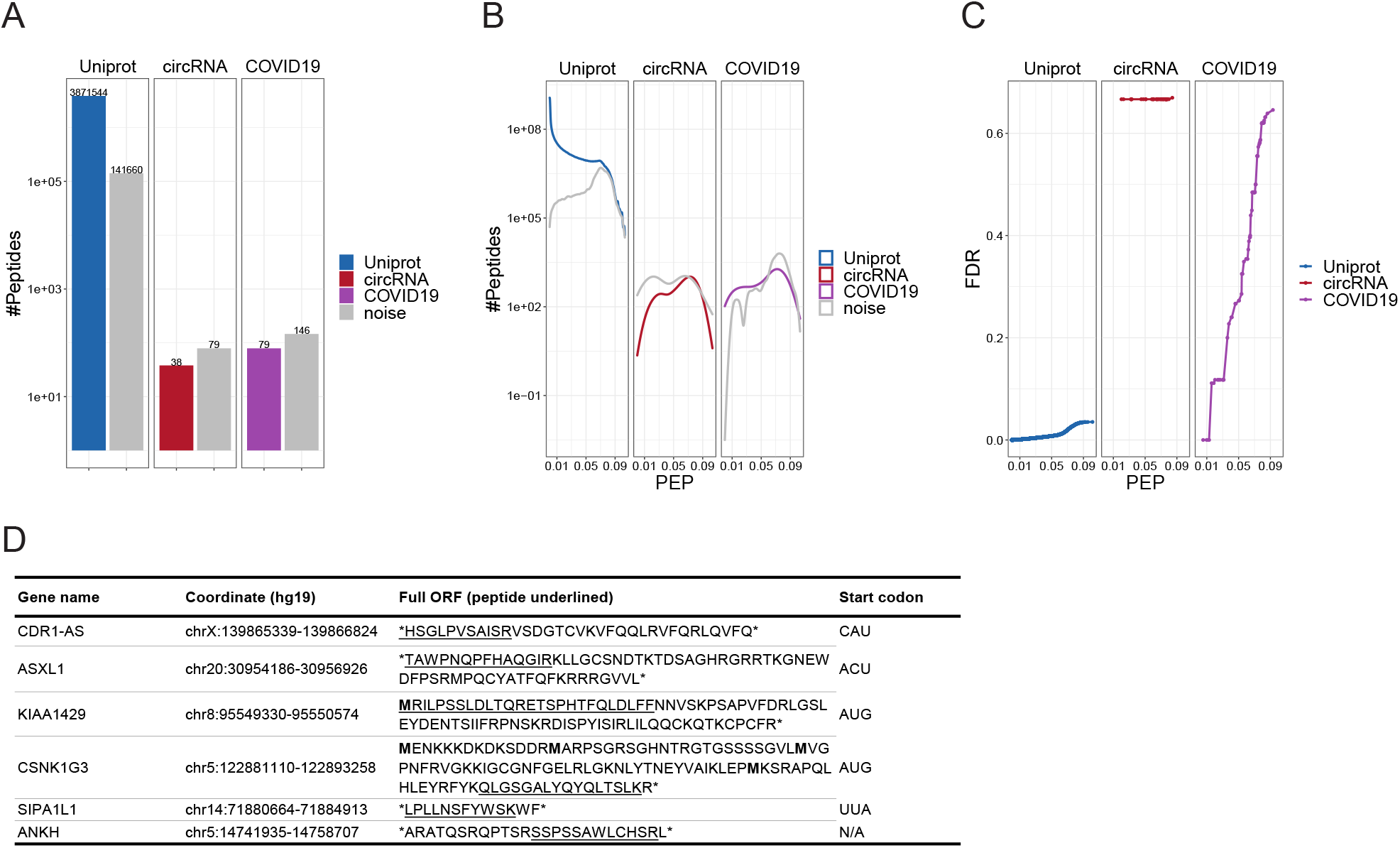
Mass spectrometry analysis. **A)** Number of target peptides and associated decoys (grey) identified from the annotated uniprot proteome (blue), the predicted ORFs across the 40 circRNAs as analysed by Heesch et al (red), and the annotated COVID19 proteome (purple). **B)** The distribution of posterior error probabilities (PEP) for target peptides from Uniprot (blue), circRNAs (red) and COVID19 (purple) and their associated decoys (noise, grey). **C)** Recalculation of FDR values based on the target-decoy strategy for each subgroup (Uniprot, circRNAs and COVID19) individually plotted as a function of PEP values. **D)** Table showing the previously published peptides from van Heesch *et al*, their predicted ORFs and predicted start codons.

Consistently, to emphasize the need for proper FDR control, three of the six BSJ-spanning peptides identified by van Heesch *et al* are preceded by stop-codons and thus translation initiation must occur cap-independently on CAU, ACU and UUA, respectively, if peptides are assumed bona fide (Fig 3D).

Collectively, this shows that without proper data analysis, anything is detectable by mass-spectrometry and additional care should be taken when conducting these experiments or when encountering them in published literature

## Conclusion

As outlined here, studying circRNA translation comes with severe pitfalls, and without meticulous data analysis, noise is easily interpreted as evidence for circRNAs translation. In contrast, exerting an unreasonable level of stringency may on the other hand result in false negatives potentially failing to disclose game changing scientific discoveries. While there may not be a golden path towards a suitable level of required stringency for each research question, it is critical to transparently estimate the contribution of noise – because there is always noise. For mass-spec, the target-decoy strategy may itself not always be sufficient for signal-noise estimates, as this procedure assumes similar score distributions for decoys and false discoveries, which in itself is controversial [24,25]. Therefore, this strategy may inflate signal and thus be more prone to false positive than negatives, and thus counterfactual analyses may provide important additional measures to ensure sound conclusions.

Evidence of circRNAs translation is based on detecting the predicted circRNAs-specific output and showing association between circRNAs and translating ribosome. As discussed above, the circRNA specific ORFs may not necessarily adhere to the host gene, and therefore any possible reading-frame across the BSJ are possible in theory. When searching for all possible scenarios, as done previously [22], it is crucial to ascertain the validity of the ORF, i.e. the presence of a canonical (preferably AUG) initiation codon. Here, for van Heesch *et al*, three of six identified peptides lack initiation codons (NUG) all together. Moreover, translation of the cap-less structure of circRNAs implies that cap-independent processes facilitates the translation. Therefore, a coherent account of circRNA translation implies the usage of endogenous IRES elements, an already controversial subject-matter not discussed further here.

CircRNAs are typically less abundant that their linear counterparts, and assumingly, if circRNA translation is true, then cap-independent initiation may also be less effective than canonical cap-dependent translation. Collectively, if at all, circRNAs are likely translated at very low levels or subjected to fast decay and as a result the derived peptides may be borderline indistinguishable from noise. Therefore, alternative sources of evidence should be considered to support or dismantle the hypothesis. Many high abundant circRNAs show cross-species conservation, and assuming that the postulated functional output from a circRNA is coupled to its biological relevance, then some level of selection pressure to preserve this output should be evident in sequence conservation analyses. Therefore, functional ORFs from conserved circRNAs species are likely to show CDS-like codon-conservation pattern, which may serve as independent evidence.

CircRNA translation is an extremely interesting hypothesis. This would not only disclose an additional proteome layer in most animals, but also shed important light on the functional relevance of potentially thousands of circRNAs. While the thrill of discovering a non-canonical game-changer in gene regulation or proteome complexity is very desirable, it is our scientific duty to uphold a high level of critical data assessment before spending valuable time and money on studying noise and artefacts. Therefore, in our ongoing scrutiny of the functional properties of circRNAs, it would likely benefit the entire circRNA research field to address our future challenges with increased stringency and additional rigor.

## Methods

### RiboSeq analysis

RiboSeq analysis was based on the procedure and data described in Stagsted *et al* [11]. Briefly, for phasing estimates and visualization, trimmed RiboSeq reads were mapped specifically to *β-ACTIN* and *GAPDH* mRNA or across circRNAs BSJ using bowtie (v1.2.3) with no mismatch tolerance (*bowtie -S -a -v* 0). Only P-sites from −8 to +6 relative to the BSJ and reads without single-mismatch alignments in the mRNA reference were considered circRNAs specific. Then, for each read-length, p-site offsets and associated quality scores were determined based on best performance on the reference CDSs (Supplementary Table 1). To assess globally the fraction of reads mapping on CDS and sno/miRNAs, reads were mapped onto hg19 using STAR (v2.7.3a) using default settings and each read-length was extracted and quantified using gencode annotations (v28lift37) with featureCounts (v2.0.0, using options `--minOverlap 0 --nonOverlap 0`) individually. Then, RiboSeq reads mapping specifically on annotated CDS and sno/miRNA regions (not overlapping any CDS) were counted.

### Mass-spec analysis

The raw mass-spec datasets where downloaded from PRIDE accession PXD006675. Maxquant (v1.6.2.10) was used for peptide identification using the complete Uniprot (UP000005640, downloaded November 10^th^, 2020), all ORFs across the 40 RiboSeq-identified circRNAs published by van Heesch *et al* [22], and the COVID19 proteome (UP000464024, downloaded November 10^th^, 2020). The Maxquant evidence files were merged and only unique peptides (targets and decoys) were kept (Supplementary Table 2).

## Availability

Data and scripts used in the analysis are available on GitHub: https://github.com/ncrnalab/circRNA_translation

## Acknowledgements

I would like to thank Prof. Gunter Meister for commenting on the manuscript, the circRtrain ITN network, and the Novo Nordisk Foundation (NNF16OC0019874) for funding.

## References

[1] X. Li, L. Yang, L.L. Chen, The Biogenesis, Functions, and Challenges of Circular RNAs, Mol. Cell. (2018). https://doi.org/10.1016/j.molcel.2018.06.034.

[2] L.S. Kristensen, M.S. Andersen, L.V.W. Stagsted, K.K. Ebbesen, T.B. Hansen, J. Kjems, The biogenesis, biology and characterization of circular RNAs, Nat. Rev. Genet. (2019). https://doi.org/10.1038/s41576-019-0158-7.

[3] A. Rybak-Wolf, C. Stottmeister, P. Glažar, M. Jens, N. Pino, M. Hanan, M. Behm, O. Bartok, R. Ashwal-Fluss, M. Herzog, L. Schreyer, P. Papavasileiou, A. Ivanov, M. Öhman, D. Refojo, S. Kadener, N. Rajewsky, P. Glazar, M. Jens, N. Pino, S. Giusti, M. Hanan, M. Behm, O. Bartok, R. Ashwal-Fluss, M. Herzog, L. Schreyer, P. Papavasileiou, A. Ivanov, M. Ohman, D. Refojo, S. Kadener, N. Rajewsky, Circular RNAs in the Mammalian Brain Are Highly Abundant, Conserved, and Dynamically Expressed, Mol. Cell. 58 (2015) 870–885. https://doi.org/10.1016/j.molcel.2015.03.027 [doi].

[4] P.L. Wang, Y. Bao, M.C. Yee, S.P. Barrett, G.J. Hogan, M.N. Olsen, J.R. Dinneny, P.O. Brown, J. Salzman, Circular RNA is expressed across the eukaryotic tree of life, PLoS One. 9 (2014) e90859. https://doi.org/10.1371/journal.pone.0090859 [doi].

[5] T.B. Hansen, T.I. Jensen, B.H. Clausen, J.B. Bramsen, B. Finsen, C.K. Damgaard, J. Kjems, Natural RNA circles function as efficient microRNA sponges, Nature. 495 (2013) 384–388. https://doi.org/10.1038/nature11993.

[6] S. Memczak, M. Jens, A. Elefsinioti, F. Torti, J. Krueger, A. Rybak, L. Maier, S.D. Mackowiak, L.H. Gregersen, M. Munschauer, A. Loewer, U. Ziebold, M. Landthaler, C. Kocks, F. le Noble, N. Rajewsky, Circular RNAs are a large class of animal RNAs with regulatory potency, Nature. 495 (2013) 333–338. https://doi.org/10.1038/nature11928; 10.1038/nature11928.

[7] M. Piwecka, P. Glažar, L.R. Hernandez-Miranda, S. Memczak, S.A. Wolf, A. Rybak-Wolf, A. Filipchyk, F. Klironomos, C.A. Cerda Jara, P. Fenske, T. Trimbuch, V. Zywitza, M. Plass, L. Schreyer, S. Ayoub, C. Kocks, R. Kühn, C. Rosenmund, C. Birchmeier, N. Rajewsky, Loss of a mammalian circular RNA locus causes miRNA deregulation and affects brain function, Science (80-.). (2017) eaam8526. https://doi.org/10.1126/science.aam8526.

[8] M. Jens, N. Rajewsky, Competition between target sites of regulators shapes post-transcriptional gene regulation, Nat. Rev. Genet. 16 (2015) 113–126. https://doi.org/10.1038/nrg3853.

[9] R. Denzler, V. Agarwal, J. Stefano, D.P. Bartel, M. Stoffel, Assessing the ceRNA Hypothesis with Quantitative Measurements of miRNA and Target Abundance, Mol. Cell. 54 (2014) 766–776. https://doi.org/10.1016/j.molcel.2014.03.045.

[10] J.U. Guo, V. Agarwal, H. Guo, D.P. Bartel, Expanded identification and characterization of mammalian circular RNAs, Genome Biol. 15 (2014) 409-014–0409-z. https://doi.org/10.1186/s13059-014-0409-z [doi].

[11] L.V. Stagsted, K.M. Nielsen, I. Daugaard, T.B. Hansen, Noncoding AUG circRNAs constitute an abundant and conserved subclass of circles, Life Sci. Alliance. 2 (2019) e201900398. https://doi.org/10.26508/lsa.201900398.

[12] I. Legnini, G. Di Timoteo, F. Rossi, M. Morlando, F. Briganti, O. Sthandier, A. Fatica, T. Santini, A. Andronache, M. Wade, P. Laneve, N. Rajewsky, I. Bozzoni, Circ-ZNF609 Is a Circular RNA that Can Be Translated and Functions in Myogenesis, Mol. Cell. 66 (2017) 22–37.e9. https://doi.org/10.1016/j.molcel.2017.02.017.

[13] N.R. Pamudurti, O. Bartok, M. Jens, R. Ashwal-Fluss, C. Stottmeister, L. Ruhe, M. Hanan, E. Wyler, D. Perez-Hernandez, E. Ramberger, S. Shenzis, M. Samson, G. Dittmar, M. Landthaler, M. Chekulaeva, N. Rajewsky, S. Kadener, Translation of CircRNAs, Mol. Cell. 66 (2017) 9–21.e7. https://doi.org/10.1016/j.molcel.2017.02.021.

[14] Y. Yang, X. Fan, M. Mao, X. Song, P. Wu, Y. Zhang, Y. Jin, Y. Yang, L.-L. Chen, Y. Wang, C.C. Wong, X. Xiao, Z. Wang, Extensive translation of circular RNAs driven by N6-methyladenosine, Cell Res. 27 (2017) 626–641. https://doi.org/10.1038/cr.2017.31.

[15] H. Ho-Xuan, P. Glažar, C. Latini, K. Heizler, J. Haase, R. Hett, M. Anders, F. Weichmann, A. Bruckmann, D. Van den Berg, S. Hüttelmaier, N. Rajewsky, C. Hackl, G. Meister, Comprehensive analysis of translation from overexpressed circular RNAs reveals pervasive translation from linear transcripts, Nucleic Acids Res. (2020). https://doi.org/10.1093/nar/gkaa704.

[16] H. Chassé, S. Boulben, V. Costache, P. Cormier, J. Morales, Analysis of translation using polysome profiling, Nucleic Acids Res. (2017). https://doi.org/10.1093/nar/gkw907.

[17] E. Sanz, J.C. Bean, D.P. Carey, A. Quintana, G.S. McKnight, RiboTag: Ribosomal Tagging Strategy to Analyze Cell-Type-Specific mRNA Expression In Vivo, Curr. Protoc. Neurosci. (2019). https://doi.org/10.1002/cpns.77.

[18] N.T. Ingolia, S. Ghaemmaghami, J.R.S. Newman, J.S. Weissman, Genome-wide analysis in vivo of translation with nucleotide resolution using ribosome profiling, Science (80-.). 324 (2009) 218–223. https://doi.org/10.1126/science.1168978.

[19] A.A. Bazzini, T.G. Johnstone, R. Christiano, S.D. MacKowiak, B. Obermayer, E.S. Fleming, C.E. Vejnar, M.T. Lee, N. Rajewsky, T.C. Walther, A.J. Giraldez, Identification of small ORFs in vertebrates using ribosome footprinting and evolutionary conservation, EMBO J. 33 (2014) 981–993. https://doi.org/10.1002/embj.201488411.

[20] J.E. Elias, S.P. Gygi, Target-decoy search strategy for increased confidence in large-scale protein identifications by mass spectrometry, Nat. Methods. (2007). https://doi.org/10.1038/nmeth1019.

[21] L. Käll, J.D. Storey, M.J. MacCoss, W.S. Noble, Posterior error probabilities and false discovery rates: Two sides of the same coin, J. Proteome Res. (2008). https://doi.org/10.1021/pr700739d.

[22] S. van Heesch, F. Witte, V. Schneider-Lunitz, J.F. Schulz, E. Adami, A.B. Faber, M. Kirchner, H. Maatz, S. Blachut, C.L. Sandmann, M. Kanda, C.L. Worth, S. Schafer, L. Calviello, R. Merriott, G. Patone, O. Hummel, E. Wyler, B. Obermayer, M.B. Mücke, E.L. Lindberg, F. Trnka, S. Memczak, M. Schilling, L.E. Felkin, P.J.R. Barton, N.M. Quaife, K. Vanezis, S. Diecke, M. Mukai, N. Mah, S.J. Oh, A. Kurtz, C. Schramm, D. Schwinge, M. Sebode, M. Harakalova, F.W. Asselbergs, A. Vink, R.A. de Weger, S. Viswanathan, A.A. Widjaja, A. Gärtner-Rommel, H. Milting, C. dos Remedios, C. Knosalla, P. Mertins, M. Landthaler, M. Vingron, W.A. Linke, J.G. Seidman, C.E. Seidman, N. Rajewsky, U. Ohler, S.A. Cook, N. Hubner, The Translational Landscape of the Human Heart, Cell. (2019). https://doi.org/10.1016/j.cell.2019.05.010.

[23] S. Doll, M. Dreßen, P.E. Geyer, D.N. Itzhak, C. Braun, S.A. Doppler, F. Meier, M.A. Deutsch, H. Lahm, R. Lange, M. Krane, M. Mann, Region and cell-type resolved quantitative proteomic map of the human heart, Nat. Commun. (2017). https://doi.org/10.1038/s41467-017-01747-2.

[24] Y. Danilova, A. Voronkova, P. Sulimov, A. Kertész-Farkas, Bias in False Discovery Rate Estimation in Mass-Spectrometry-Based Peptide Identification, J. Proteome Res. (2019). https://doi.org/10.1021/acs.jproteome.8b00991.

[25] N. Gupta, N. Bandeira, U. Keich, P.A. Pevzner, Target-decoy approach and false discovery rate: When things may go wrong, J. Am. Soc. Mass Spectrom. (2011). https://doi.org/10.1007/s13361-011-0139-3.

